# Flocculation of a cyanobacterium confers defense against bacterial predation

**DOI:** 10.1101/2025.10.27.684795

**Authors:** Shylaja N. Mohandass, Alice C.Z. Collins, Fabian D. Conradi, Luke P. Allsopp, Conrad W. Mullineaux

## Abstract

Many cyanobacteria, including unicellular species, are capable of flocculation: the formation of floating linked assemblages of many thousands of cells. Flocculation is a highly regulated process requiring both type IV pilus activity and the production of extracellular polysaccharides. Under standard laboratory conditions flocculation often slows culture growth, and its physiological advantages remain unclear. Proposed benefits include self-shading as protection from excess light exposure and flotation in the water column. Here, we determine whether flocculation can serve as a method of defense against bacterial predation. Using the unicellular cyanobacterium *Synechocystis* sp. PCC 6803, we show that flocculation is strongly triggered by exposure to live cells of “foreign” bacteria, including the opportunistic pathogen *Pseudomonas aeruginosa*. The established *P. aeruginosa* virulence arsenal includes bacterial warfare systems for competition for resources and acquisition of nutrients by way of interbacterial competition. Here, we establish the use of these strategies for direct predation, via the use of Type VI Secretion Systems. Comparisons of *P. aeruginosa* co-cultures with either wild type *Synechocystis* or a non-flocculating mutant revealed that *Synechocystis* flocculation minimizes both growth of *P. aeruginosa* and cell lysis of *Synechocystis*. This in turn reduces the impact of *P. aeruginosa* on *Synechocystis* growth by mechanistically limiting the photosynthetic products that *P. aeruginosa* can access. From these data, we propose that the type VI secretion system of *P. aeruginosa* can be used for predation and that the primary function of cyanobacterial flocculation is for defense against microbial predation.

**IMPORTANCE:** Bacteria in the environment rarely, if ever, grow in isolation. Some phenotypic traits can only be understood in the context of interspecies interactions. Here, we examine interactions between two bacterial species with very different modes of metabolism. *Synechocystis* sp. PCC 6803 is a freshwater cyanobacterium that grows photosynthetically using light, water, CO_2_ and minerals. *Pseudomonas aeruginosa* is a soil bacterium and an opportunistic pathogen that depends on a supply of organic molecules for survival. When *P. aeruginosa* is co-cultured with *Synechocystis* in mineral medium, it lyses *Synechocystis* cells, using its Type VI secretion system to access nutrients via predation. The presence of *P. aeruginosa* (and other foreign bacteria) triggers a *Synechocystis* response in which it aggregates to form floating assemblages or flocs. Floc formation creates a defensive barrier to *P. aeruginosa* predation, and we suggest this complex behavior is best understood as a defense against microbial predation in the wild.

## INTRODUCTION

Cyanobacteria are globally important as primary producers and are found in a wide variety of marine, freshwater and terrestrial habitats (1). Much research on cyanobacteria has focused on their oxygenic photosynthesis and associated autotrophic metabolism, however cyanobacteria have other fascinating aspects to their biology. Filamentous cyanobacteria provide examples of sophisticated prokaryotic multicellularity (2), and unicellular cyanobacteria are capable of complex and co-operative behaviors (3). Co-operative behavior is exhibited by the unicellular freshwater cyanobacterium *Synechocystis* sp. PCC 6803 (hereafter *Synechocystis*), which under the appropriate conditions forms large floating assemblages or ‘flocs’, which can contain thousands of cells (3–7). *Synechocystis* floc formation requires both the type IV pilus (T4P) apparatus (6, 7) and the production of a specific sulfated exopolysaccharide (EPS) termed Synechan (4). Floc formation is a tightly regulated process which is promoted by environmental factors including nutrient limitation (6), and blue light acting via the cyanobacteriochrome photoreceptor Cph2 (7).

*Synechocystis* cells in the center of dense flocs show signs of nutrient stress (7), and comparison of wild type (WT) *Synechocystis* to a non-flocculating mutant shows that flocculation slows growth under standard laboratory conditions (7). This hints at some unknown benefit of flocculation. One possibility is that flocculation promotes self-shading and reduces exposure to excessive or harmful light. In the thermophilic cyanobacterium *Thermosynechococcus vulcanus*, flocculation is induced by low temperatures and blue light (8) to promote self-shading, reducing exposure to excessive or harmful light. This requires production of extracellular cellulose stimulated by a network of photoreceptors acting through the intracellular second messenger cyclic di-GMP (9, 10). In *Synechocystis*, flocculation has been suggested to benefit cells by trapping gas bubbles for flotation, which might allow flocculated cells to find a more favorable level in the water column (4). Additionally, flocculation has been proposed to promote formation of mutualistic bacterial communities. *Synechocystis* was previously shown to form more durable biofilms in the presence of a heterotrophic partner, *Pseudomonas taiwanensis* (11). This enhanced stability suggested a mutualistic relationship in which *P. taiwanensis* aerobic metabolism prevents the excessive build-up of oxygen in the biofilm (11), which may be extrapolated to floating flocs (3).

Here, we test another possibility, that cyanobacterial flocculation serves primarily as a defense mechanism against predation. We selected the opportunistic pathogen *Pseudomonas aeruginosa* as a model bacterial predator as it actively engages in bacterial warfare (12–14). *P. aeruginosa* lyses competitor cells using an arsenal of weapons, including contact-independent cell lysis molecules such as bacteriocins, pyocins and porins (12, 15), and contact-dependent systems such as the type VI secretion system (T6SS) (12–14). Bacterial warfare studies typically focus on heterotrophic species and are considered a competition for resources (13, 14, 16). However, these antibacterial weapons could be employed for direct predation. As *P. aeruginosa* is dependent on organic molecules for survival, predation of the phototrophic *Synechocystis* may be significant in co-culture in mineral medium. Here, we studied how *P. aeruginosa* interacts with *Synechocystis* when co-cultured in mineral medium. We show predation on *Synechocystis* cells by *P. aeruginosa* is required for predator success, and is predominantly accredited to the action of the T6SSs. In response *Synechocystis* employs flocculation as a defense mechanism against *P. aeruginosa* predation.

## RESULTS

### Flocculation of *Synechocystis* is triggered by exposure to foreign bacteria

The cyanobacterium *Synechocystis* sp. PCC 6803 has evolved in laboratory cultures into a range of substrains with different phenotypic characteristics (17). Here we used the PCC-M substrain (18), which is highly motile on surfaces and readily flocculates in planktonic cultures (7). We induced and quantified flocculation in *Synechocystis* using a method previously outlined (7), in which planktonic cultures are grown using BG11 medium in 6-well plates with gentle shaking, where flocculation was recorded after 48h. BG11 mineral medium supports photosynthetic growth but lacks organic carbon sources (19). To quantify flocculation, we recorded standardized digital camera images of the wells and measured the normalized standard deviation in the image pixel values, defining a numerical aggregation score. A high standard deviation indicates an inhomogeneous cellular distribution within the well and therefore a high level of flocculation or aggregation of the cells (7). Our control, a Δ*hfq* mutant, lacks cell appendages (20) and does not flocculate (7). As expected, these aggregation scores corroborate WT flocculation under our conditions, whilst Δ*hfq* cells remained evenly dispersed and consequently give much lower aggregation values (Fig 1 A,B). Upon addition of foreign bacterial cells to these cultures, enhanced flocculation of *Synechocystis* was observed (Fig 1C). Interestingly, using either the potentially aggressive *Pseudomonas aeruginosa* PA14 (21) or the non-pathogenic standard laboratory strain *Escherichia coli* Top10 - widely used as a benign prey strain in inter-microbial competition assays (22) - flocculation appears independent of foreign cell genus and intrinsic behavior. Both species induced similar very dense flocs (Fig 1C) which was reflected in the change in aggregation scores observed when compared to WT *Synechocystis* monocultures (Fig 1D). To check if the enhanced flocculation was simply a cell density-driven response, we supplemented the wells with a similar number of extra *Synechocystis* cells, however this had no effect (Fig 1C,D). Next, we used calcofluor white staining (23) to investigate EPS production in *Synechocystis* flocs. Flocs from *Synechocystis* monoculture generate some EPS but have a relatively open structure (Fig 2A). However, exposure to *P. aeruginosa* induced increased production of EPS (Fig 2B), which appeared to form a protective barrier at the floc edge (Fig 2C).

**Figure 1:**
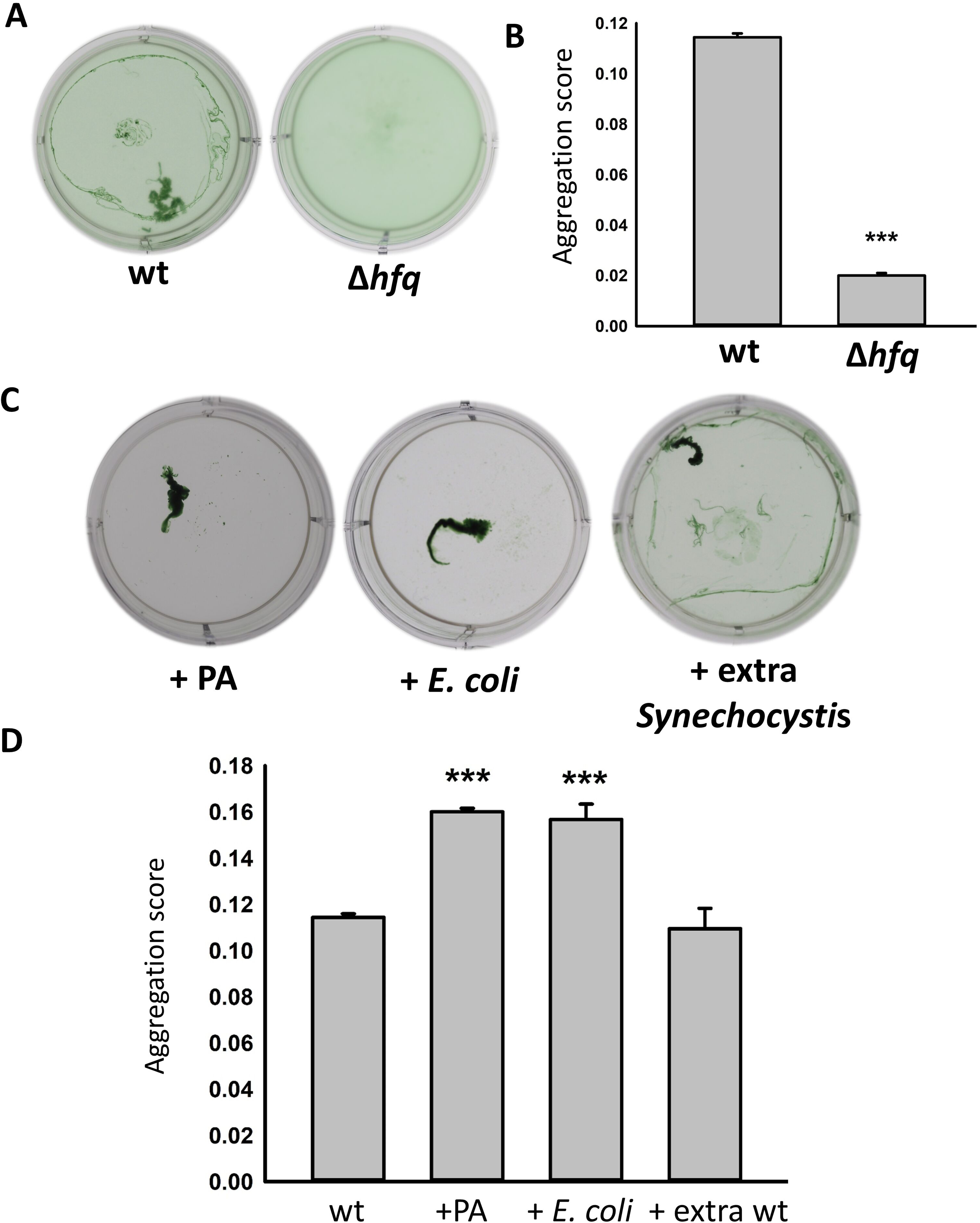
*Synechocystis* flocculation is triggered by exposure to foreign bacteria. **A.** Representative flocculation assays for pure cultures of *Synechocystis* wild type and Δ*hfq*. **B.** Mean aggregation values for pure cultures of wild type and Δ*hfq*. **C.** Representative flocculation assays for *Synechocystis* wild type co-cultured with *P. aeruginosa* PA14, Top10 *E. coli* or with the addition of an equivalent amount of extra *Synechocystis.* D. Mean aggregation values for flocculation assays of the co-cultures. All measurements are means from 3 biological replicates. Error bars indicate SE and *** indicates significant difference from pure wild type *Synechocystis* (*p* < 0.0001).

**Figure 2.**
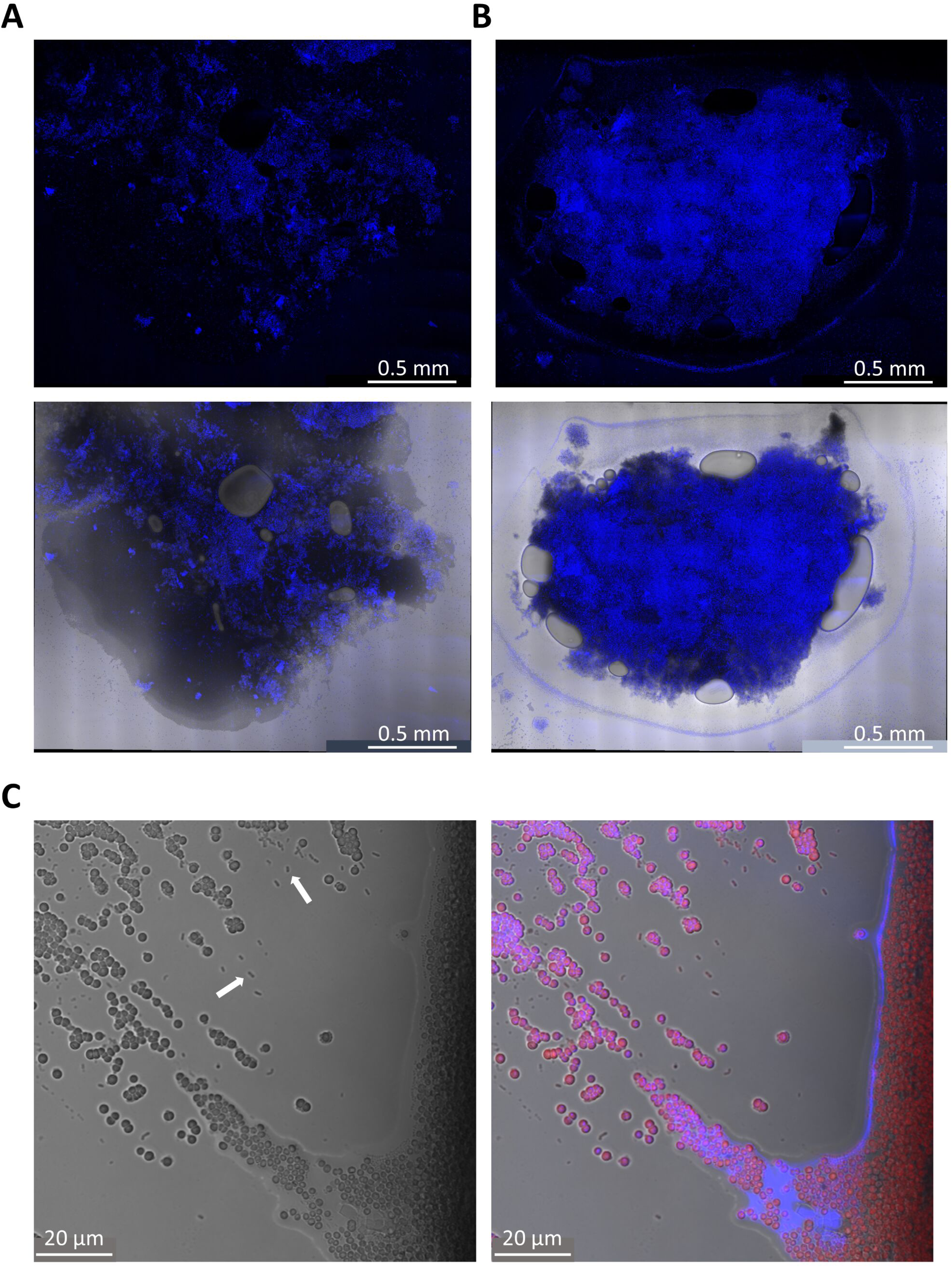
Dense *Synechocystis* flocs are associated with high concentrations of extracellular polysaccharide. **A,B.** Wide-angle microscopic images of floc samples showing Calcofluor white fluorescence (in blue) merged with brightfield (grayscale) in the lower images. **A.** Floc from a *Synechocystis* pure culture **B.** Floc from a *Synechocystis* co-culture with *P. aeruginosa* PA14. **C.** Higher-magnification image of the edge of a floc from a *Synechocystis*:*P. aeruginosa* co-culture. Brightfield image on the left: *P. aeruginosa* cells are visible as small rods, with examples highlighted by the arrows. On the right is a merged image with brightfield plus chlorophyll fluorescence (red) and Calcofluor white fluorescence (blue).

### *Synechocystis* flocculation is triggered by exposure to live or heat-killed *P. aeruginosa* cells

To explore potential factors responsible for triggering denser *Synechocystis* flocculation, we tested the effects of filtered media from *P. aeruginosa* PA14 monocultures and *Synechocystis*:PA14 co-cultures upon flocculation after 48 h. No significant increase in aggregation scores was observed (Fig 3). Since there is evidence that kin cell lysis can serve as a bacterial danger signal (24), we next added mechanically-broken *Synechocystis* cells, however this had only a marginal effect on flocculation (Fig 3). Additionally, an extract of PA14 EPS failed to induce additional flocculation (Fig 3). However, addition of intact heat-killed PA14 cells did significantly elevate aggregation scores with a visible effect (Fig 3), although these flocs appeared less compact and dense than those induced by live PA14 cells (Figs 1,3). Taken together, the results suggest that *Synechocystis* flocculation is induced by contact with foreign bacterial cell components independent of species (Fig 1). The greater response to live *P. aeruginosa* cells (Fig 1) compared to heat-killed cells (Fig 3) may be due to cell proliferation during the 48h incubation (see Fig 4) and/or be due to *P. aeruginosa* specifically targeting *Synechocystis* through an active mechanism (25).

**Figure 3.**
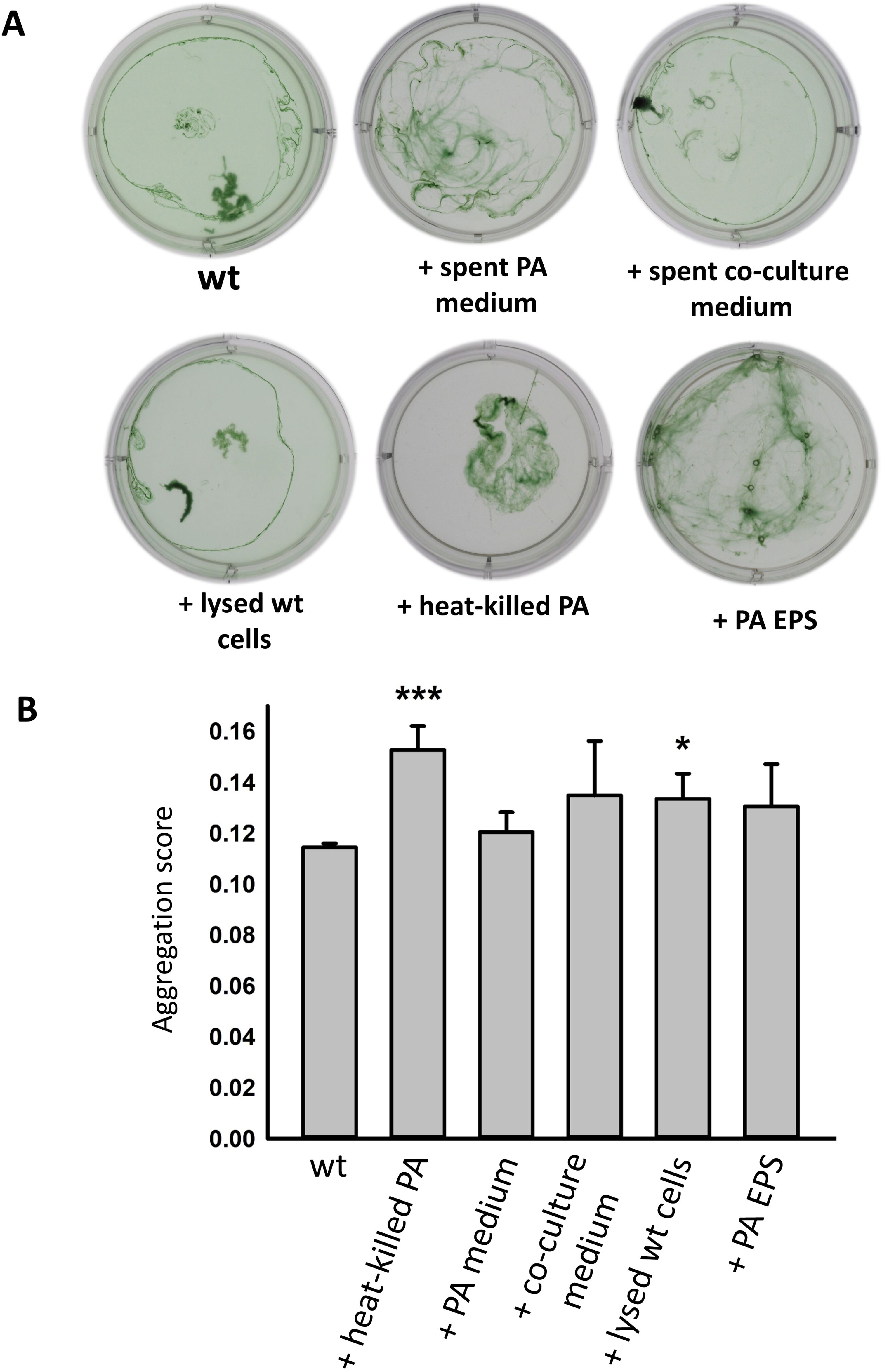
Effects of culture extracts on *Synechocystis* flocculation. **A.** Representative flocculation assays for *Synechocystis* wild type incubated with heat-killed *P. aeruginosa* cells, filtered medium from *P. aeruginosa* cultures and *P. aeruginosa*:*Synechocystis* co-cultures, mechanically-lysed *Synechocystis* cells and *P. aeruginosa* EPS extracts. **B.** Mean aggregation values from these conditions. Error bars indicate SEM from 3 biological replicates. Asterisks indicate significant difference from culture without additions (***: *p* < 0.0001; *: *p* < 0.05).

**Figure 4.**
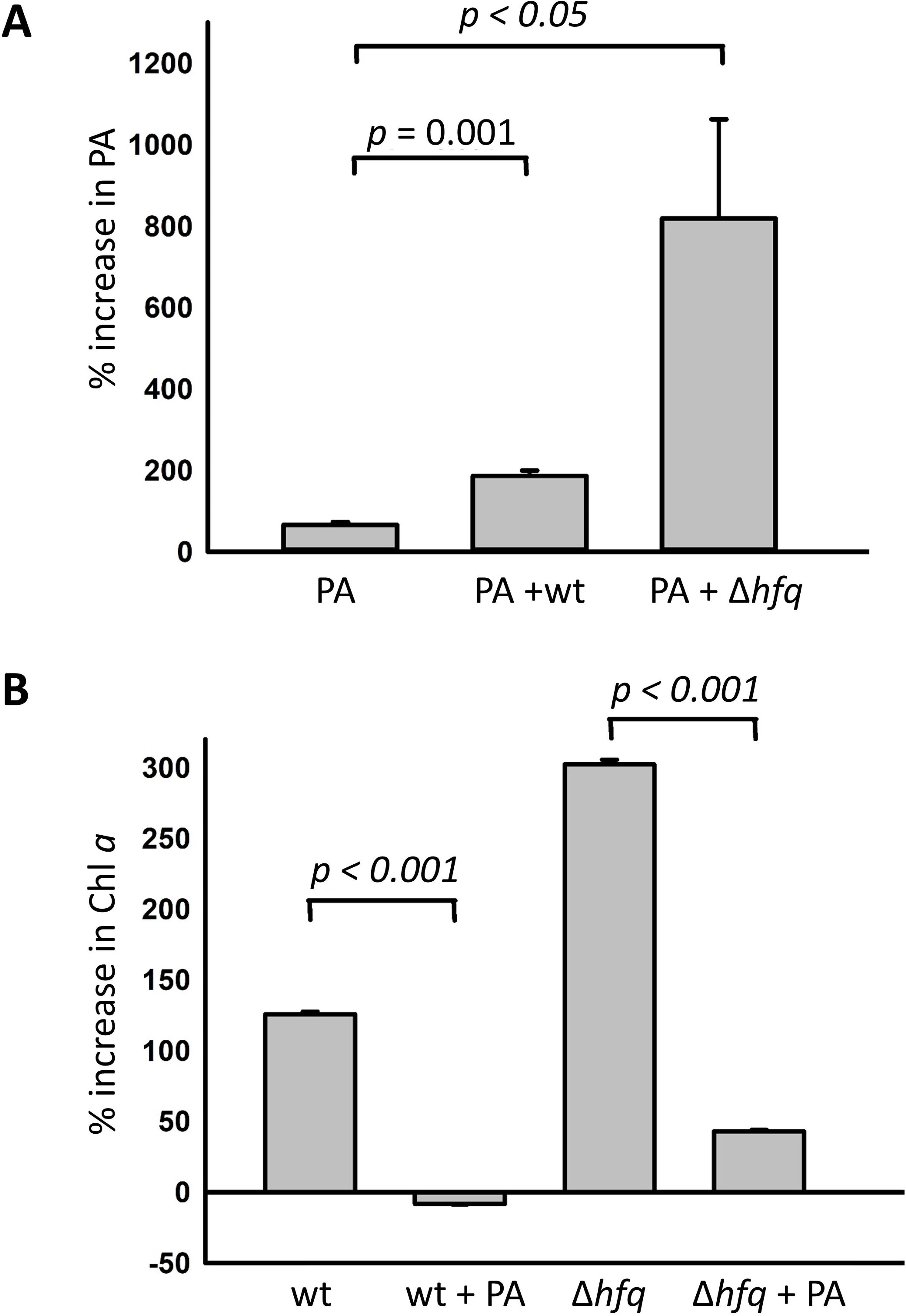
Growth of *Synechocystis* and *P. aeruginosa* in co-culture for 48h. **A**. Growth of *P. aeruginosa* in BG11 medium in pure culture or together with *Synechocystis* wild type or Δ*hfq.* Growth assessed from CFU counts. **B.** Growth of *Synechocystis* strains in pure culture or together with *P.* aeruginosa, assessed from chlorophyll concentration. All values are means from 3 replicates. Error bars indicate SEM and *p*-values from t-tests are shown.

### Flocculation of *Synechocystis* reduces the growth of *P. aeruginosa* in co-culture

To assess the impact of *Synechocystis* and *P. aeruginosa* on their respective growth during co-culture, we quantified growth of the two species over 48h in BG11 medium. *P. aeruginosa* growth was assessed by counting colony-forming units (CFU) from the co-cultures on LB-agar plates, on which *P. aeruginosa* grows rapidly but *Synechocystis* does not. As expected, PA14 cell growth in monoculture was minimal as BG11 medium lacks organic carbon sources (Fig 4A). To assess the additional impact of *Synechocystis* flocculation on growth, we compared PA14 co-cultures using either WT *Synechocystis* or the non-flocculating Δ*hfq* mutant (7). Here, a significant increase in PA14 growth occurred in WT *Synechocystis* co-culture, but PA14 growth was greater in co-culture with Δ*hfq* (Fig 4A). This indicates that photosynthetic *Synechocystis* products can be exploited for *P. aeruginosa* growth, and that these products are more readily available from the non-flocculating Δ*hfq* strain.

As assessing *Synechocystis* growth via CFU is difficult due to cell adherence and the slow appearance of colonies, we extracted chlorophyll from the co-cultures as a proxy for *Synechocystis* cell density. Comparison of WT *Synechocystis* with Δ*hfq* mutant monocultures is complicated, as Δ*hfq* grows faster than wild type under our culture conditions (Fig 4B), likely because flocculation reduces the efficiency of light absorption and mineral supply to cells in the center of flocs (7). Nevertheless, chlorophyll concentration at 48h shows that the presence of PA14 greatly impacts *Synechocystis* (Fig 4B), where the population deficit is significantly higher for Δ*hfq* than the WT (Fig 4B). This is consistent with the faster proliferation of PA14 in co-culture with Δ*hfq* (Fig 4A).

The impacts of co-culture on both *P. aeruginosa* and *Synechocystis* growth (Fig 4B) suggest that *P. aeruginosa* may be actively predating *Synechocystis* cells for survival rather than just utilizing secreted photosynthetic products. Microscopic examination of the co-cultures (Fig 5 A,C,E) highlighted the presence of lysed *Synechocystis* cells, which were rarely observed in pure cultures (Fig 5 B,D). A significant proportion of the *Synechocystis* cells become indistinct in brightfield and show much lower chlorophyll fluorescence in co-culture (Fig 5 C,E), indicating cell lysis and subsequent loss or degradation of photosynthetic pigments. *P. aeruginosa* cells are observable as much smaller, non-fluorescent rods, and are often seen clustering around *Synechocystis* cells, hinting at the occurrence of a contact-dependent interaction (Fig 5 C,E).

**Figure 5.**
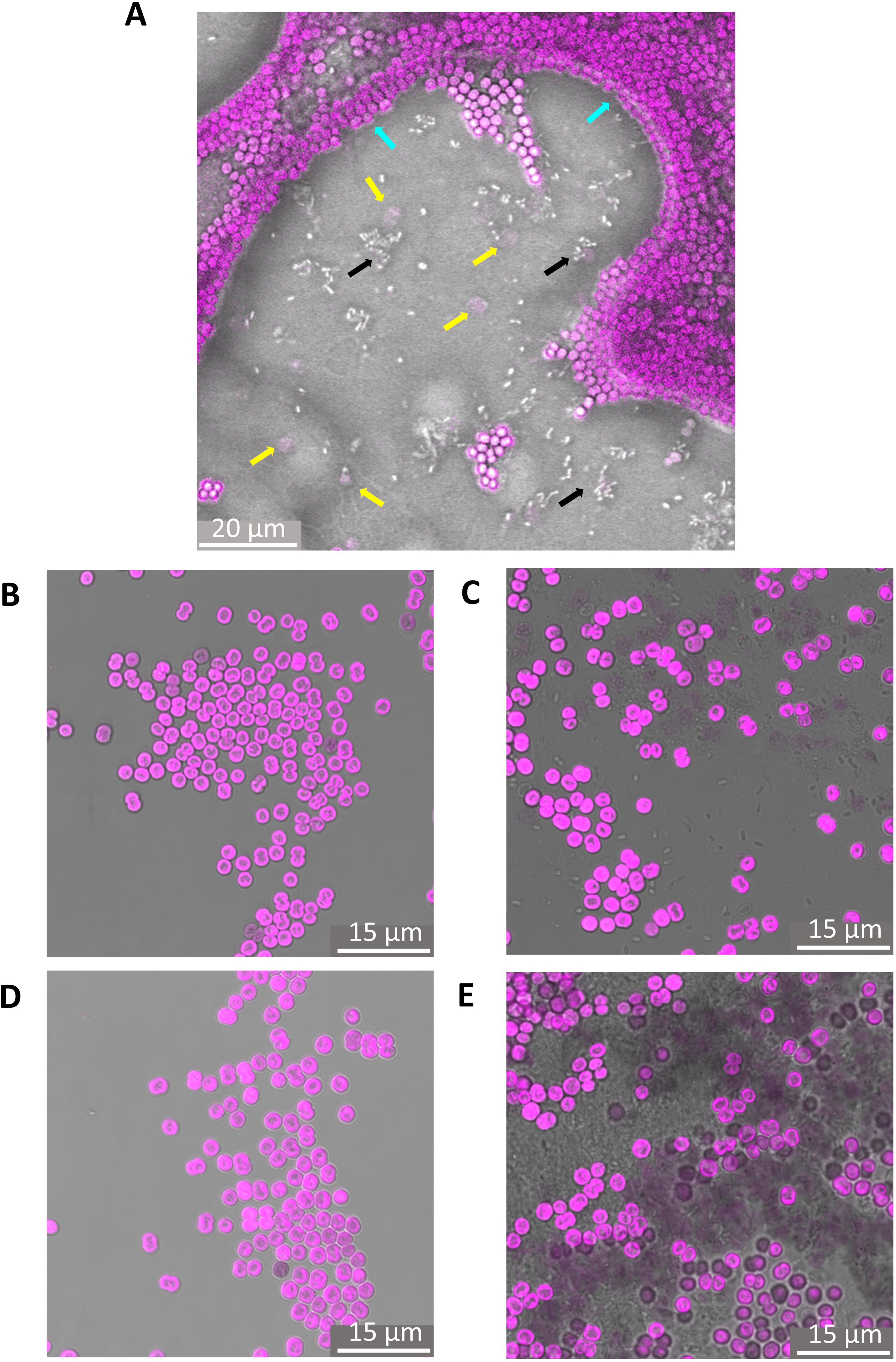
Addition *P. aeruginosa* to *Synechocystis* cultures results in cell death. Chlorophyll fluorescence from *Synechocystis* cells is shown in magenta, merged with brightfield images in grayscale. **A.** The edge of a *Synechocystis* floc (wild type plus *P. aeruginosa*). Note *Synechocystis* cells (eg cyan arrow) *P. aeruginosa* cells (eg black arrows) and some very pale *Synechocystis* cells that appear to have been lysed (eg yellow arrows). **B-E.** representative images from cultures and co-cultures. **B.** *Synechocystis* wild type. **C.** Wild type plus *P. aeruginosa*. **D.** *Synechocystis* Δ*hfq.* **E.** Δ*hfq* plus *P. aeruginosa*. Note the presence of *P. aeruginosa* cells and compromised *Synechocystis* cells with indistinct edges and minimal fluorescence in C. and E.

### *P. aeruginosa* uses multiple mechanisms to lyse *Synechocystis* cells

To explore the mechanisms that *P. aeruginosa* may use to lyse *Synechocystis* cells, we set up co-cultures with *P. aeruginosa* PA14 mutants deficient in either contact-dependent or contact-independent antibacterial weapons. The newly-constructed PA14 H123^-^ lacks all three of the *P. aeruginosa* T6SS (H1-, H2- and H3-T6SS (12)), whilst PA14 *prtN::tn* (26) lacks the positive regulator required for pyocin production (27). Both mutants strongly impacted *Synechocystis* growth in co-culture (Fig 6A), and microscopic examination indicated noticeable *Synechocystis* cell death induced by both mutants (Fig 6B-E). The pronounced effects seen with the PA14 H123^-^ mutant indicate that a contact-independent mechanism can induce *Synechocystis* cell death. In line with this, we found that filtered *P. aeruginosa* culture media triggered the appearance of large numbers of “ghost” *Synechocystis* cells with minimal chlorophyll fluorescence (Fig 6F): such cells were rarely seen in healthy *Synechocystis* cultures (Fig 5 B,D).

**Figure 6.**
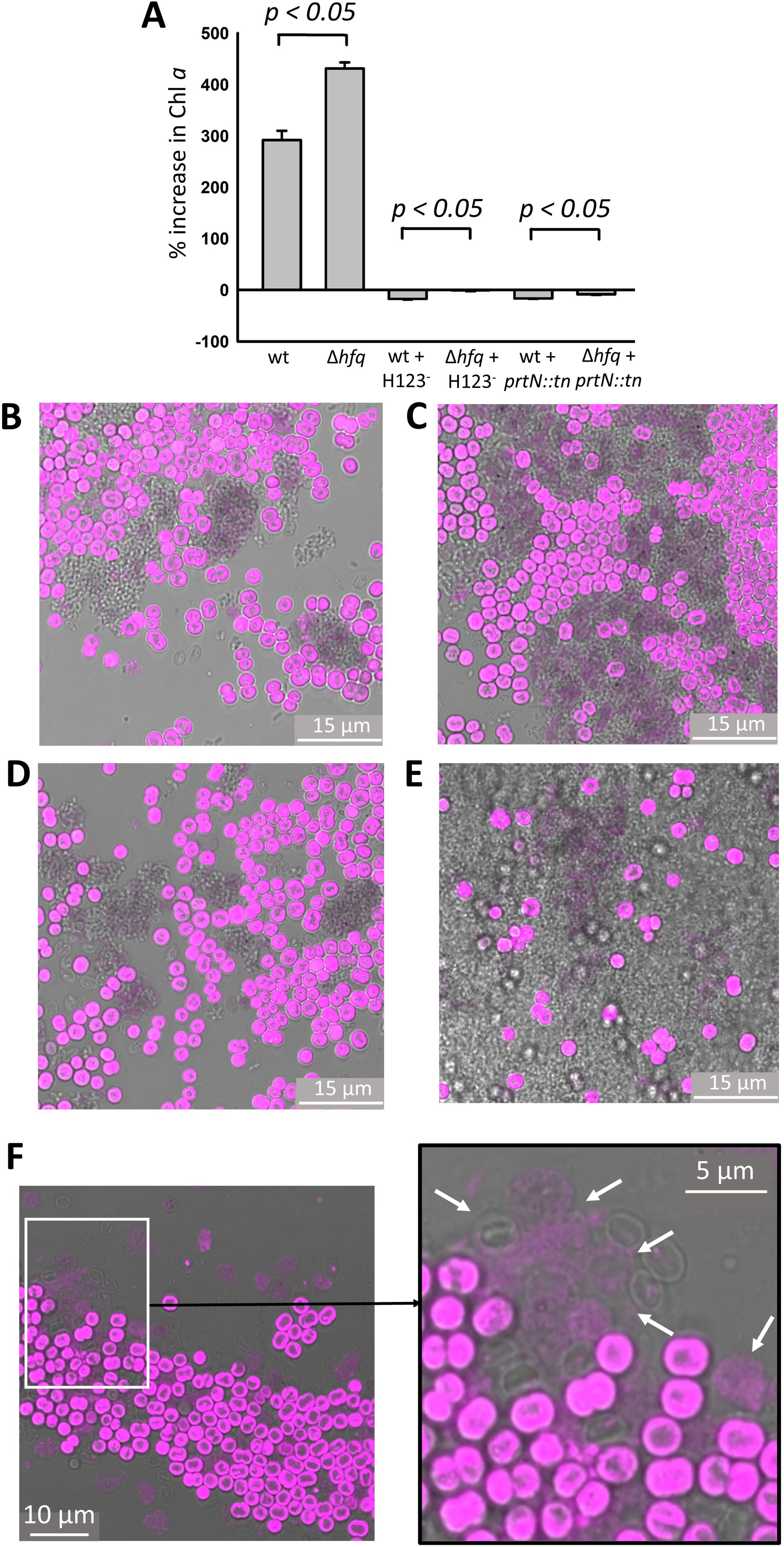
Effects of *P. aeruginosa* mutants and spent medium on *Synechocystis* strains. **A.** Growth of *Synechocystis* wild type and Δ*hfq* in co-culture with *P. aeruginosa mutants*, assessed from chlorophyll concentration after 48 h. Means from 3 biological replicates, error bars indicate SEM. **B-E.** Fluorescence micrographs showing *Synechocystis* cell death induced by the PA14 mutants. Chlorophyll fluorescence in magenta merged with brightfield (grayscale). **B.** Wild type *Synechocystis* with PA14 H123^-^. **C.** *Synechocystis* Δ*hfq* with PA14 H123^-^. **D.** Wild type *Synechocystis* with PA14 *prtN::tn*. **E.** *Synechocystis* Δ*hfq* with PA14 *prtN::tn*. **F.** Fluorescence micrograph, with detail, showing *Synechocystis* cells compromised by exposure to filtered *P. aeruginosa* medium. Arrows highlight *Synechocystis* cells in various stages of collapse.

To comprehensively examine the effects of *P. aeruginosa* PA14 WT and mutant cells on *Synechocystis*, fluorescence micrographs of *Synechocystis* cells from the co-cultures were investigated to visualize the native chlorophyll fluorescence from the thylakoid membranes. Healthy cells are roughly spherical in shape. The thylakoid membrane layers in these cells are rather irregular, but their distal surfaces are quite smooth because they are contained within the smooth cell envelope layers. In cells from *P. aeruginosa* co-cultures, a proportion of cells had convoluted distal surfaces to their thylakoid membranes, suggesting disruption of the cell envelope. We quantified this effect by measuring the perimeter:area ratio, and found significant increases in the mean ratio for WT *Synechocystis* with WT PA14 co-culture, and for *Synechocystis* Δ*hfq* with either WT, H123^-^ or *prtN::tn* PA14 co-culture (Fig 7). The presence of these disrupted cells indicates direct cell attack that may be mediated in either contact-dependent or contact-independent fashion.

**Figure 7:**
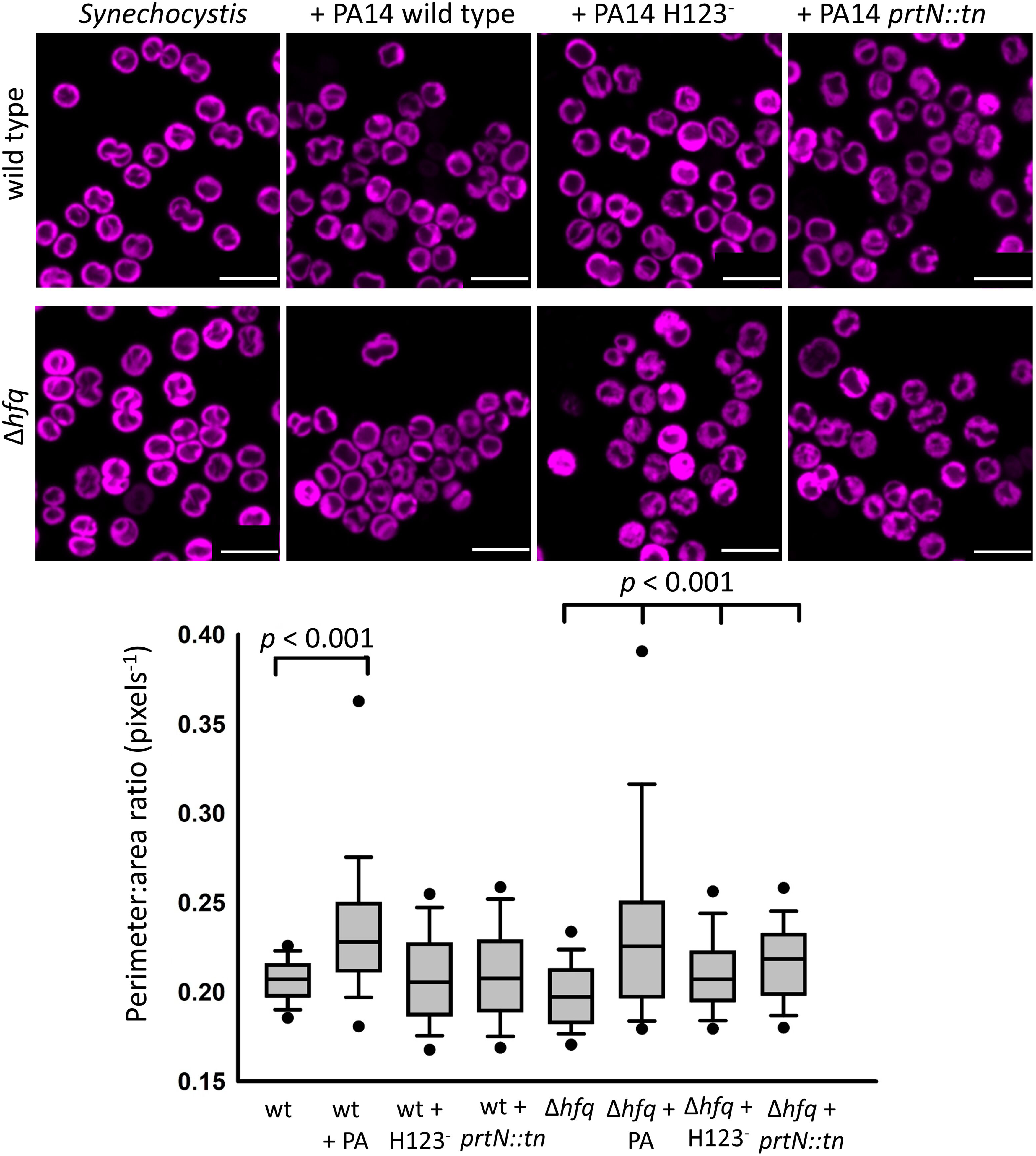
*Synechocystis* cell structure changes induced by exposure to *P. aeruginosa* wild type and mutants. Specimen fluorescence micrographs are shown at the top (chlorophyll fluorescence in magenta; all scalebars 5 µm) and quantitation of the cell perimeter: area ratio is shown below (*n =* 100 cells in each case) with standard box and whisker plots. *p* values from t-tests comparing the appropriate *Synechocystis* strain in pure culture vs co-culture are indicated.

### Contact-dependent weapons are more effective for supporting growth of *P. aeruginosa* in co-culture

Although the two PA14 mutants were similarly effective in killing *Synechocystis* cells (Fig 6,7), there were striking differences in their growth in co-culture with *Synechocystis* strains (Fig. 8). As expected, both mutants failed to grow in BG11 medium in the absence of *Synechocystis* (Fig 8 A,B). In the presence of *Synechocystis* WT or Δ*hfq*, PA14 H123^-^ showed only very modest growth after 48h (Fig 8A), however PA14 *prtN::tn* proliferated strongly, especially in co-culture with *Synechocystis* Δ*hfq* (Fig 8B). In this situation, growth of PA14 *prtN::tn* (Fig 8B) was even faster than growth of WT *P. aeruginosa* (Fig 4A). The strong negative impact of the H123^-^ mutation on PA14 growth in *Synechocystis* co-culture (Fig 8) demonstrates the importance of the contact-dependent mechanism (T6SS) for PA14 growth in this condition and confirms that PA14 deploys both contact-dependent and contact-independent weapons against *Synechocystis*.

**Figure 8.**
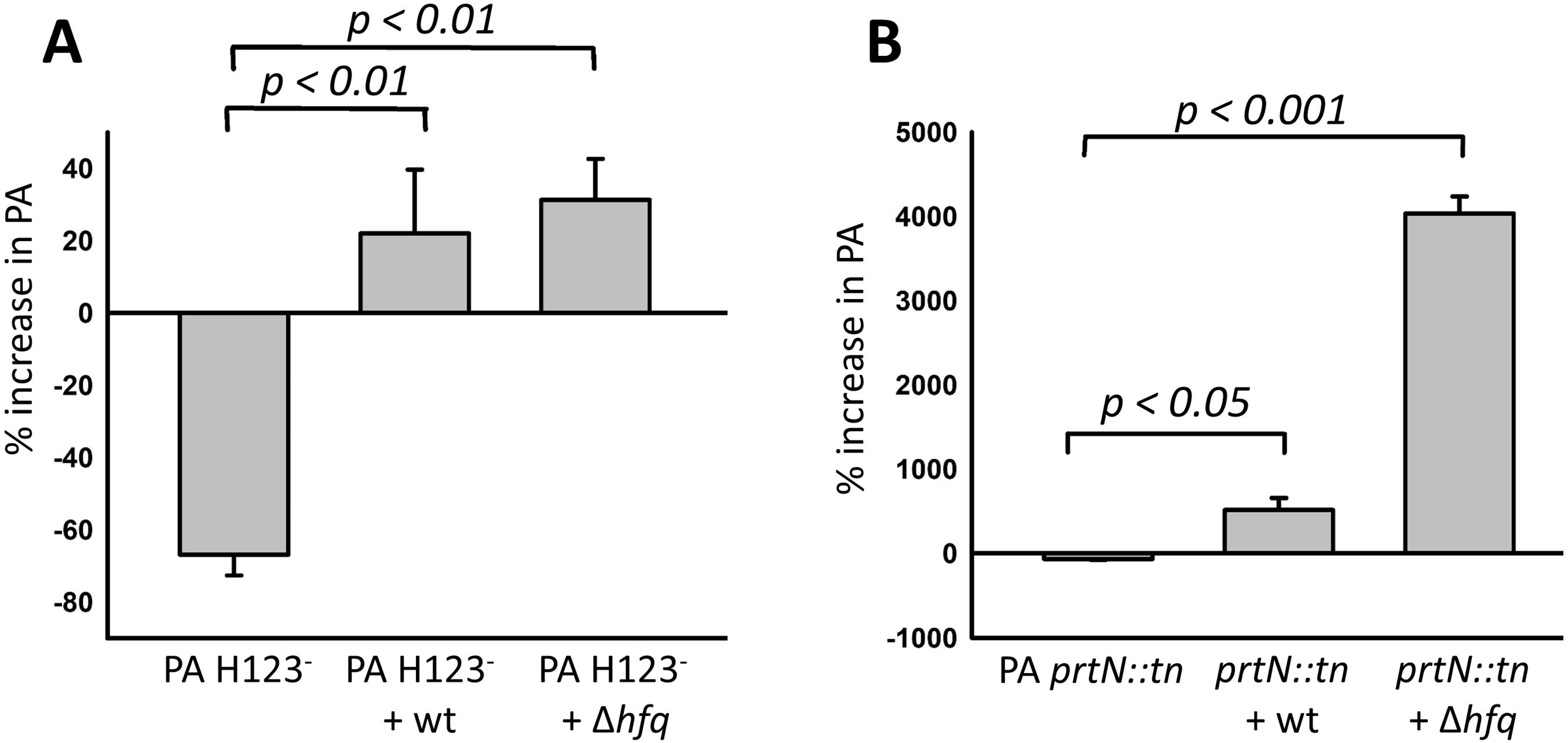
Growth of *P. aeruginosa* mutants in co-culture with *Synechocystis* strains. Growth was assessed from CFU counts after 48 h of co-culture. Data are means from 3 biological replicates and error bars indicate SEM. **A.** PA H123^-^, lacking T6SS. **B.** PA *prtN::tn*, deficient in production of pyocins. Differences between growth of PA *prtN::tn* and PA H123^-^ are significant: *p* = 0.026 for comparison of the PA strains in co-culture with *Synechocystis* wt; *p* = 0.0004 for comparison of the PA strains in co-culture with *Synechocystis* Δ*hfq*. Note that proliferation of PA H123^-^ is much slower than proliferation of PA *prtN::tn* in this condition.

Taken together, our results indicate that both contact-independent attack with pyocins and contact-dependent attack with T6SS are similarly effective at killing *Synechocystis* cells. However, *P. aeruginosa* cell proliferation is greatly impaired in the PA14 H123^-^ mutant lacking T6SS, whilst PA14 *prtN::tn*, which retains the T6SS but not pyocins, proliferates strongly in co-culture with *Synechocystis*. This indicates that *P. aeruginosa* requires its T6SS to fully benefit from predation.

## DISCUSSION

### *P. aeruginosa* employs its Type VI secretion systems for predation on *Synechocystis*

Predation amongst bacteria is well established, with examples ranging from the intracellular parasitism of *Bdellovibrio* (28) to bacterial hunting by swarms of *Myxococcus xanthus* (29). *P. aeruginosa* is highly motile and equipped with a formidable array of antibacterial weapons (12) often used to eliminate competitors (14). Here, we show that *P. aeruginosa* can employ these weapons for direct predation, enabling proliferation in co-culture with a cyanobacterium in mineral medium, where organic compounds are only available as photosynthate from the cyanobacterium. Our results with *P. aeruginosa* mutants (Figs 6,7) and filtered *P. aeruginosa* medium (Fig 6) indicate that *P. aeruginosa* can lyse *Synechocystis* cells by both contact-dependent and contact-independent mechanisms. Contact-independent mechanisms appear similarly effective for lysing *Synechocystis* cells but not for PA14 survival. This is exemplified in Fig 8, as a mutant lacking pyocins for contact-independent cell lysis thrives in *Synechocystis* co-culture, whereas the H123-mutant lacking T6SS survives, but grows only very slowly. T6SS are contact-dependent weapons that operate by direct injection of toxic proteins into target cells, leading to cell death (13, 30). The H2-T6SS and H3-T6SS of *P. aeruginosa* have also been shown to be involved in metal iron acquisition with specific effectors that target iron, copper and molybdate (31–34). As the T6SS directly injects toxic bacterial-derived proteins into target cells, the advantage of this contact-dependent system is that target cell lysis releases cell contents into the milieu for immediate consumption by the adjacent attacking cell. By contrast, remote lysis by contact-independent systems releases cell contents which become diluted in the surrounding milieu. Our micrographs of *P. aeruginosa*:*Synechocystis* co-cultures support close contact between lysed *Synechocystis* cells and predating *P. aeruginosa* (Fig 5), but further work is required to understand the interaction dynamics. T6SSs are effective “disintegration weapons” through their delivery of lytic toxins (35), and it is possible that this may serve as a particularly effective way to release prey cell contents for consumption. However, recent work has elegantly shown that killing by the T6SS using effector sets that result in slow lysis results in a graded release of nutrients allowing for more efficient use by the attacking organism (16), and this may be the case with predation on *Synechocystis*.

### Direct contact with foreign bacteria induces the formation of dense flocs of *Synechocystis*

Several studies have shown that *Synechocystis* cells assemble into flocs in appropriate conditions in pure planktonic cultures (4, 6, 7). Here we have shown that exposure to foreign bacteria induces the formation of dense *Synechocystis* flocs (Fig 1) exhibiting enhanced extracellular polysaccharide production (Fig 2). This dense flocculation is induced to similar extents by both the benign *Escherichia coli* Top10 and the more aggressive *Pseudomonas aeruginosa* PA14 (Fig 1). Dense flocculation does not appear to be a specific response to attack, since can be partially induced by heat-killed PA14 cells but is not induced by exposure to filtered PA14 medium (Fig 3), although this medium is effective in killing some *Synechocystis* cells (Fig 6F). Our results suggest that dense flocculation is triggered by contact with the cell surfaces of foreign bacteria. It is likely that discrimination between “self” and “non-self” is involved, because flocculation is not a simple response to cell density: as the addition of an equivalent number of extra *Synechocystis* cells does not trigger denser flocculation (Fig 1). The method by which *Synechocystis* senses contact with foreign bacteria is unknown, but it is likely that T4P are involved. Although the requirement for floc formation of T4P (6, 7), and production of a sulfated EPS is established (4), it is not clear whether T4P are required to trigger the response by forming initial cell-cell contacts, or by promoting EPS production, or both. T4P are involved in surface sensing in many bacteria (36), and *Synechocystis* is known to respond to surface contact with changes in gene expression (37). In addition, *Synechocystis* encodes an extensive and diverse set of T4P minor pilins which may recognize multiple targets (38, 39). As a working hypothesis, we suggest that *Synechocystis* pilus tips adhere to components found on foreign cell envelopes, and the resulting tension in the pilus triggers signal transduction via cyclic di-GMP (40) that leads to EPS production and dense floc formation (7, 9). Previously*, P. taiwanensis* was shown to stabilize co-culture biofilms with *Synechocystis* (11) and another cyanobacterium (41). The mechanism is unclear, but was suggested to involve regulation of oxygen levels in the biofilms by *P. taiwanensis* (11). In our experiments with planktonic cells and shaken culture wells with an airspace, O_2_ concentration is unlikely to be a significant variable, due to ready atmospheric equilibration. Nevertheless, the presence of heterotrophic bacteria strongly induces *Synechocystis* cell aggregation. We suggest that direct contact sensing of *P. taiwanensis* in the biofilms may explain the previously observed stabilizing effect.

### Flocculation in *Synechocystis* provides a defensive barrier against bacterial predation

Dense *Synechocystis* flocs are associated with high concentrations of extracellular polysaccharide, and microscopic examination of the edges of flocs suggest that they could provide an effective physical barrier to penetration of contact-dependent and independent mechanisms employed by bacteria such as *P. aeruginosa* (Figs 2, 5A). Production of EPS capsules has been shown in another bacterium to be an effective defense against attack via T6SS (42). Here we show that *P. aeruginosa* proliferation in *Synechocystis* co-culture is greater with the non-flocculating *Synechocystis* Δ*hfq* mutant than with the WT (Figs 4,8), providing strong evidence that flocculation reduces predation. We cannot exclude the possibility of other roles of flocculation such as flotation (11) or shading (5,7), and indeed the roles of photosensors in promoting cell aggregation in *Synechocystis* (7) and other cyanobacteria (9, 10) suggest a role in modulating the light environment. However, encounters with predatory microbes are common in the environment. Amoebae and other protozoa are known to graze on cyanobacteria (43–45), and our laboratory experiments suggest that predation by heterotrophic bacteria may also be commonplace. We argue that the primary and most pressing function of cyanobacterial flocs may be to provide protection from microbial predation.

## MATERIALS AND METHODS

### Bacterial strains and growth conditions

*Synechocystis* strains used were the motile PCC-M substrain of *Synechocystis* sp PCC 6803 (18) and the Δ*hfq* mutant constructed in the same background (20). *Synechocystis* starter cultures were grown in BG11 medium (19) supplemented with 2-{[1,3-Dihydroxy-2-(hydroxymethyl)propan-2-yl]amino}ethane-1-sulfoniacid (TES) buffer (pH 8.2) at 30°C in plastic tissue culture flasks (Sarstedt), under continuous white light illumination (∼15 μmol photons m^-2^ s^-1^) with shaking (120 rpm). Cultures were also maintained on BG11 plates containing 1.5% (w/v) Bacto-agar (VWR) supplemented with TES buffer (pH 8.2) and 0.3% (w/v) sodium thiosulfate.

The heterotrophic bacteria (*Pseudomonas aeruginosa* strains and *Escherichia coli* Top10 from Thermo-Fisher Scientific) were cultured in LB liquid medium and on LB agar plates (Thermo-Fisher Scientific) at 37°C with appropriate antibiotics. All *Pseudomonas aeruginosa* strains were in the PA14 background (21), including the mutants *prtN::tn* (26) and the newly-constructed H123^-^.

### Construction of the PA H123^-^ mutant

Genomic, PCR and plasmid vector DNA was purified using Qiagen DNeasy Blood and Tissue, QIAquick PCR Purifications kit and QIAprep Spin Miniprep kits, respectively. DNA fragments were amplified with either KOD Hot Start DNA Polymerase (Merck) or standard Taq polymerase (NEB) as described by the manufacturer with the inclusion of Betaine (Sigma) or DMSO (Sigma). pKNGrsmA restore was generated by amplifying the WT copy of *rsmA* using primers P1 and P2 (Table S1) and cloning this into pKNG101 using ApaI and SmaI. Restriction endonucleases and ligase were used according to the manufacturer’s specifications (NEB). All constructs were DNA sequenced and confirmed to be correct before use by Eurofins Genomics. Plasmids were introduced into *E. coli* strains by heat shock transformation and to *P. aeruginosa* strains by conjugation. Strain PA14 H123^-^ was engineered in this study by restoring the wild-type copy of *rsmA* in the PA14*rsmA* H123^-^ (22) background (*rsmA*, H1-T6SS (*tagQ1-vgrG1b*), H2-T6SS (*tssA2-clpV2*) and H3-T6SS (*tssB3-clpV3*)).

### Flocculation assays

Freshly cultured *Synechocystis* cells were pelleted by centrifugation at 4000 x g for 5 min and resuspended in fresh BG11 medium (without TES buffer, as this is a potential organic food source for heterotrophs) to an OD_750_ of 0.5 (Jenway 6300 spectrophotometer). Cell density was ∼2.8 x 10^8^ cells ml^-1^ as assessed by counting in a hemocytometer (Neubauer counting chamber). The resuspended *Synechocystis* cultures were transferred into Corning® Costar® TC-treated 6-well plates (Sigma-Aldrich) with 5 ml of culture per well. Where appropriate for coculture assays, overnight cultures of the heterotrophic bacteria *(P. aeruginosa*, *E. coli*) were standardized to OD_600_ of 1.5 (equating to a cell density of ∼9 x 10^8^ ml^-1^ for *P. aeruginosa*) and a 50 μl aliquot was added to each well. For flocculation assays with additional *Synechocystis* cells, a *Synechocystis* culture was concentrated to ∼9 x 10^8^ cells ml^-1^ and a 50 µl aliquot added to the well. The 6-well plates were then sealed and incubated at 30°C for 48 h with white light illumination at 30 μmol photons m^−2^ s^−1^ with shaking at 75 rpm (16-mm orbit diameter/2-mm stroke in an SI50 orbital incubator; Stuart Scientific, UK). All assays were performed in triplicate. After 48 h growth, the 6-well plates were removed from the incubator, placed on a light box (Medalight LP-300N cold cathode fluorescent lamp) and imaged using using an Olympus OM-D (E-M1 Mark II) camera vertically positioned on a stand (Kaiser 205361) at a height of 40 cm. All images were taken with standard specifications (ISO 200, shutter speed 40, F5.6, default white balance). Olympus Capture software was used for positioning and focusing. Images were analyzed with Image J (46) and numerical aggregation values were calculated from the normalized standard deviation of the images as in (7).

### Growth measurements

*Synechocystis* and *P. aeruginosa* cell densities in co-culture were assessed at the start of the experiment and after 48 h. The contents of the wells were transferred to 50 ml tubes (Sarstedt) and vigorously shaken and vortexed to disperse the flocs and ensure even distribution of the cells. *P. aeruginosa* cells in the co-culture were then quantified by colony-forming unit (CFU) assays. Coculture samples were serially diluted 10-fold in BG11 medium (without TES buffer) and 10 μl aliquots were spotted onto *Pseudomonas* isolation agar (Millipore, Sigma-Aldrich) followed by incubation overnight at 35°C and colony counting.

Growth of *Synechocystis* in co-culture was assessed by measuring chlorophyll *a* as a proxy for cell density. Chlorophyll in 1 ml aliquots was extracted with methanol and the concentration calculated from absorbance at 665 nm as previously described (47).

### Extracts from cultures

A variety of extracts from cell cultures were tested for their influence on *Synechocystis* flocculation. In all these cases, flocculation was assessed after 24-28 h. Spent *P. aeruginosa* medium was obtained by centrifuging aliquots of overnight grown *P. aeruginosa* cultures (OD_600_ 1.5) at 10,000 x g for 10 min. The supernatant was then filter sterilized using a 0.2-μm syringe filter (Fisher Scientific). Fifty micro litres of filtered spent medium were added to the well for the assay. For the spent medium from co-cultures, *P. aeruginosa*-*Synechocystis* co-cultures were grown as described above for flocculation assays. Cells were then pelleted by centrifugation, and the supernatant was filtered and added to flocculation assays as for the spent *P. aeruginosa* medium.

Extracellular polymeric substances (EPS) were extracted by ethanol precipitation according to (48, 49). Supernatants from 100 ml of freshly grown *P. aeruginosa* cultures were mixed with 95% ethanol in a 1:3 ratio (v/v) and incubated at 4°C overnight to precipitate EPS. The EPS was separated by centrifuging the mixture at 4000 x g for 20 min followed by incubation of the pellet at 37°C to evaporate the solvent. The pellet was resuspended in double-distilled water to 0.4% w:v by vigorous mixing until it completely dissolved. The solution was then passed through a 0.45 μm cellulose acetate membrane (Thermo-Fisher Scientific) to remove any remaining cells. Subsequently, a volume of 50 μl of the extracted EPS was added to the flocculation assay. *Synechocystis* cell lysates were prepared by mixing 350 μl of healthy *Synechocystis* cells at an OD_750_ of 0.5 with 150-212 µm glass beads (Sigma) at a 1:1 ratio in an Eppendorf tube. The mixture was vortexed for 10 cycles using a Disruptor Genie (Scientific Industries), with each cycle consisting of 1 min of disruption followed by a 1 min pause at 4°C. The lysate was then centrifuged at 13000 x g for 10 min and 50 μl of the supernatant was added to a flocculation assay.

Heat-killed *P. aeruginosa* cells were prepared according to (50) with some modification. Overnight cultures at an OD_600_ of 1.5 were centrifuged at 12,000 x g for 10 min and the pellet was resuspended in the same volume of BG11 medium (without TES buffer) followed by incubation at 92°C for 2 h. 50 μl of the heat-killed culture was added to a flocculation assay.

### Preparation of samples for microscopic imaging

Floc samples from the coculture assay were carefully removed using blunted pipette tips and gently pressed between two 24×60 mm microscope coverslips. The edges were sealed to keep the flocs stationary during imaging. EPS (β-polysaccharide) in the flocs was visualized using calcofluor white staining (Sigma-Aldrich). Flocs were gently transferred to a Nunc Lab-Tek 2-well chamber slide using blunt pipette tips, followed by addition of calcofluor white (300 mg l^−1^, 100 μl) (41). The slides were then incubated in the dark for 30 min at 30°C. After incubation, the flocs were carefully washed with 1X PBS buffer and placed between microscope coverslips as before.

### Confocal imaging and image analysis

All imaging was with a Leica Stellaris 8 confocal laser scanning microscope equipped with a 63X oil immersion objective (numerical aperture 1.4). Chlorophyll fluorescence from *Synechocystis* cells was imaged with excitation from a 405 nm diode laser and emission at 670-720 nm. *P. aeruginosa* was visualized in brightfield mode. Calcofluor white fluorescence was excited at 405 nm and detected at 420-490 nm. Images were recorded in 8-bit, 512×512 pixel format with 8x line averaging at 400 Hz scan speed and visualized with Leica LAS AF software. Widefield images were obtaining in tiling mode, combining sets of images recorded as before except without line averaging. Each tile in the grid was autofocused before tile scanning.

*Synechocystis* cell area and perimeter analysis were performed in ImageJ using the chlorophyll fluorescence images. Cell boundaries were determined by thresholding and individual cells in clear view were selected. Cells undergoing division were handled manually.

## ACKNOWLEDGMENTS

Research in the CWM laboratory was supported by The Leverhulme Trust (RPG-2020-054) and the Biotechnology and Biological Sciences Research Council (BB/W019698/1). We thank Kavitha Rangasamy for her contributions to experimental work. Work in the LPA laboratory is supported by the Academy of Medical Sciences/Wellcome Trust/Government Department of Business, Energy and Industrial Strategy/the British Heart Foundation/Diabetes UK Springboard Award [SBF006\1161] and by a Biotechnology and Biological Sciences Research Council grant BB/Y00048X/1. LPA has also benefited from supported from the National Institute of Health and Care Research (NIHR) through the Imperial Biomedical Research Centre (BRC) Grant. ACZC PhD studentship was funded by the NHLI.

## Supplementary Material

**Table S1:**
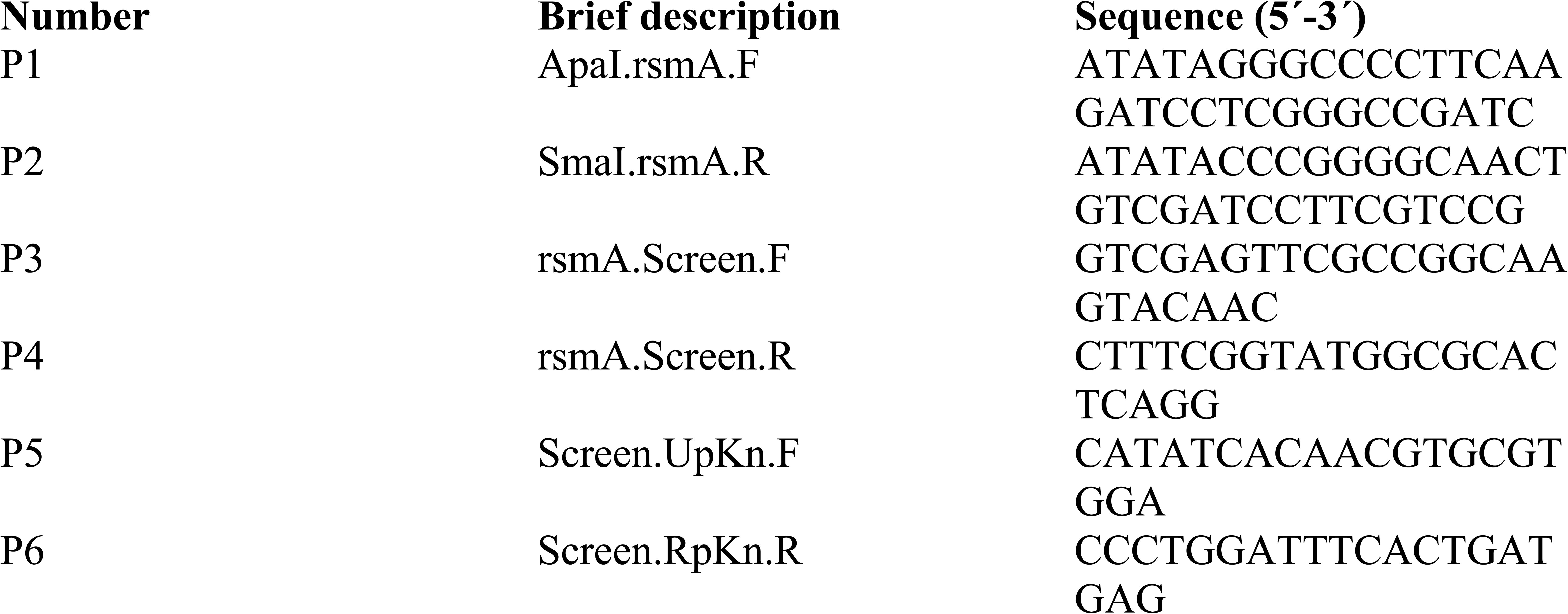
PCR primers used in generation of the PA H123^-^ mutant.

